# MRI Brain Templates of the Male Yucatan Minipig

**DOI:** 10.1101/2020.07.17.209064

**Authors:** Carly Norris, Jonathan Lisinski, Elizabeth McNeil, John W. VanMeter, Pamela VandeVord, Stephen M. LaConte

## Abstract

The pig is growing in popularity as an experimental animal because its gyrencephalic brain is similar to humans. Currently, however, there is a lack of appropriate brain templates to support functional and structural neuroimaging pipelines. The primary contribution of this work is an average volume from an iterative, non-linear registration of 70 male Yucatan minipig subjects whose ages ranged from five to seven months. In addition, several aspects of this study are unique, including the comparison of linear and non-linear template generation, the characterization of a large and homogeneous cohort, an analysis of effective resolution after averaging, and the evaluation of potential within template bias as well as a comparison with a template from another minipig species using a “left-out” validation set. We found that within our highly homogeneous cohort, non-linear registration produced better templates, but only marginally so. Although our T1-weighted data were resolution limited, we preserved effective resolution across the multi-subject average, produced templates that have high gray-white matter contrast, and demonstrated superior registration accuracy compared to the only known alternative minipig template.

## 1. Introduction

Across a broad spectrum of biomedical research, the pig is emerging as an important experimental animal that is more human-relevant than rodents while balancing the monetary and ethical costs associated with non-human primates. In neuroscience, pigs are advantageous because their gyrencephalic anatomy, developmental time course, and neurochemistry are similar to that of human brains (Yun et al., 2011; Conrad et al., 2014; Ishizu et al., 2000; Jakobsen et al., 2006; Fang et al., 2005; Lind et al., 2007). Because of this, it is not surprising that neuroimaging studies using pigs have been conducted across the full range of modalities, including PET, CT, MRI, EEG and fNIRS (Lind et al., 2007; Sauleau et al., 2009; Roura et al., 2016; Schubert et al., 2016). Despite their promise and increasing use, neuroimaging software and analysis pipelines for pigs are currently lacking compared to primates, rodents, and humans. As we outline here, one of the most critical current needs for pig neuroimaging is the development of templates for brain mapping studies.

Most available neuroimaging analysis software was designed hand-in-hand with human data. Fortunately, though, these tools are also generally very flexible. Thus to accommodate other species, many functions can be applied directly or readily adapted by establishing new default parameters such as field-of-view, resolution, whole-brain and tissue specific volumes, spatial smoothness, and hemodynamic response functions. Frequently the most data and labor intensive component of species-specific analysis pipelines is the development of appropriate templates and atlases. Though both terms are sometimes used interchangeably, we refer to a template as a reference brain that defines a standardized coordinate “brain space” and an atlas as providing the additional benefit of defined anatomical labels. Although a single subject can be used as a template, it is more common to use multi-subject data to better capture population features. Within an analysis pipeline, the template defines the reference space. For a neuroimaging study, the data can be normalized into this space through linear transformations and/or nonlinear warping operations. One of the key benefits of normalizing to a template is that it enables group statistics, with increasing statistical power for every additional subject and the potential to make inferences that apply beyond the study sample to the broader population (Mazziotta et al., 2001). Templates also provide a standard underlay image upon which to visualize multi-subject statistical results from structural and functional analyses. Lastly, templates can provide a coordinate system to report the spatial locations of those statistics.

As mentioned, normalization (registering or aligning one brain into the space of another) can be done through linear or non-linear transformations. Technically, linear transformation matrices have six degrees of freedom to enable 3D translations and rotations, while affine transforms additionally include scaling, reflections, and shear, using twelve degrees of freedom. Both are sometimes referred to as “linear,” in contrast to non-linear transformations which use thousands of parameters. Usually normalization is performed in the context of analyzing study data with a pre-existing template. It is generally believed that the greater the degrees of freedom for the normalization approach, the better the alignment (Crivello et al., 2002; Klein et al., 2009). In practice, however, a broad range of techniques often lead to relatively equivalent results, depending on the research goal and the smoothness and resolution of the study’s data in relation to the template. Computation time verses diminishing returns on alignment quality is usually a secondary consideration, but can occasionally become paramount in the context of high resolution datasets, large numbers of subjects, and long convergence times. Most relevant for the work reported here, normalization is also the primary step used to generate templates from multiple subjects. Since the motivation for multi-subject templates is to avoid being biased to any one individual’s variability, the ideal goal is to capture the population mean at every location in the brain. As outlined by Fonov et al. (2011), the methods for achieving this can be categorized as relying predominantly on feature matching or intensity matching strategies (Fonov et al., 2011). Although there are a variety of template building approaches for human data (Collins et al., 1994; Ardekani et al., 2005; Avants et al., 2006; Lorenzen & Joshi, 2003), most tend to be initiated by linear registration followed by an iterative non-linear refinement step. Many of the recent non-human templates tend to use tools from the major neuroimaging processing packages (SPM, FSL, and AFNI) and often adopt methods of early human template creation such as manual skull stripping, intensity normalization, and anterior commissure to posterior commissure (AC-PC) alignment followed by normalization techniques (Fox et al., 1985; Collins et al., 1994; Evans et al., 1992, 1993; Ullmann et al., 2015; Nitzsche et al., 2018; Conrad et al., 2014; Ella & Keller, 2015; Seidlitz et al., 2018).

The present study utilizes the Yucatan minipig (Panepinto et al., 1978) to develop a T1-weighted template. Adult commercial pig breeds can be challenging in terms of experimental protocols, equipment designs, and feed and care costs because they can weigh from 140 to 270 kg (300 to 600 lbs)(Estrada et al., 2008). Thus piglets and minipigs have become popular for overcoming this drawback. There are currently no MRI brain templates available for the Yucatan. The limited number of pig templates that do exist include the domestic pig (Saikali et al., 2010), the neonatal piglet (Conrad et al., 2014), and the Göttingen minipig (Watanabe et al., 2001). The Yucatan is known to be gentle, has an adult weight ranging from approximately 70 to 90 kg (150 to 200 lbs), and has been used extensively for developing surgical and experimental techniques as well as models for metabolic syndrome, biocompatability, skin lesions, pharmacology, toxicology, and cardiovascular disease (Estrada et al., 2008; Eubanks et al., 2006; Curtasu et al., 2019; Mattern et al., 2007; Quesson et al., 2011; Montezuma et al., 2006; Pak et al., 2006; Lin et al., 1998; Hurtig et al., 2019; Swindle et al., 1990, 2011; Witczak et al., 2006; Lopez et al., 2017). The Yucatan has also been used to develop a reliable brain tumor model for glioblastoma (Khoshnevis et al., 2017) and recent studies have utilized MRI to investigate head and neck vasculature, mechanical properties related to TBI, and stroke (Guertler et al., 2018; Habib et al., 2013; Platt et al., 2014). As a topic related to templates, it should be noted that for neurosurgical procedures, stereotaxic coordinates in the form of topological atlases for pigs have been available for decades (Lind et al., 2007; Yoshikawa, 1968; Salinas-Zeballos et al., 1986; Félix et al., 1999) and stereotaxic methodology continues to be an area of research and development (Bjarkam et al., 2009; Rosendal et al., 2010; Glud et al., 2017). Indeed, for the Yucatan, the study most closely related to this report comes from a stereotaxic comparative study between 6 animals imaged at 1.5 T and compared to their axially sectioned histology (Yun et al., 2011).

Unlike recent template development efforts in pigs and other animal models (Saikali et al., 2010; Conrad et al., 2014; Watanabe et al., 2001; Ella & Keller, 2015; Seidlitz et al., 2018; Nitzsche et al., 2018, 2015; Hikishima et al., 2011; Love et al., 2016; McLaren et al., 2009; Quallo et al., 2010), the data collected here was not specifically motivated by template creation from the onset. Rather, the T1-weighted data used in this project consisted of a single scan among several in a multimodal, multi-time point study of blast-induced traumatic brain injury. Because of the nature of the study, acquisition time was limited. This is offset, though, by the large number of subjects (70 total) and their high degree of homogeneity in terms of age and weight. Thus, for our group, the study data provided both a need (as a prerequisite to further data analysis) and an opportunity for this work. Moreover, the data used here are representative of data that other groups might collect during imaging sessions, where time is limited by the need to collect a broad number of scans while minimizing both the subject’s time under anesthesia and scanning costs.

As will be described further, this project ultimately generated four templates. Although these templates are the primary contribution, several aspects of this study are unique. First, we compared both linear and non-linear registration. Second, we used a large and homogeneous cohort compared to other currently available animal atlases. This homogeneity allowed us to characterize the spatial variance that occurs with normal genetic variation across a narrow range of subject ages and weights. Third, we have characterized the effective resolution of our templates via the spatial Fourier transform. Fourth, we evaluated our templates and compared them to the Göttingen minipig (Watanabe et al., 2001) using a “left-out” validation set to compare registration errors to anatomical regions with independent data. The four templates are publicly available (https://lacontelab.github.io/VT-Yucatan-MRI-Template/). In addition, we have archived our analysis scripts, the original T1-weighted volumes, and each subject’s estimated gray matter, white matter, and cerebral spinal fluid maps.

## 2. Materials and Methods

### 2.1. Image Acquisition

MRI data were acquired in male Yucatan minipigs in accordance with the Virginia Tech Institutional Animal Care and Use Committee. Imaging was performed during the baseline condition of a multi-time point study of traumatic brain injury. In total, 72 subjects were scanned using six cohorts, ranging from 6 to 19 subjects per cohort, collected over a 3 year period. Two subject scans were omitted due to noticeable artifacts. The remaining 70 subjects had a mean age of 5 months 16 days (minimum 4.9 months, maximum 7.3 months) and a mean weight of 23.18 kg (minimum 17.4 kg, maximum 30.3 kg). Scanning was performed with a Tim Trio 3 Tesla scanner (Siemens AG, Erlangen) using 3 elements of an 8-channel spine array coil. Subjects were supine with their heads near the foot of the table. The anesthesia and end-tidal *CO*_2_ tubing was run through a waveguide in the control room along with a fiber-optic cable for an MRI-compatible pulse oximeter, passing approximately 4.6 m from the wall to the scanner’s isocenter. T1-weighted anatomical volumes (resolution = 1 × 1 × 1 *mm*^3^; TR = 2300 *ms*; TE= 2.89 *ms*; TI = 900 *ms*; FOV = 256 *mm*^2^; FA = 8°; BW = 140 *Hz/pixel*) were collected with a three-dimensional magnetization prepared rapid acquisition gradient echo (MPRAGE) pulse sequence (Mugler & Brookeman, 1990).

### 2.2. Image Preprocessing and Template Generation

Images were processed in AFNI (Cox, 1996) using procedures adapted from recent animal brain template reports (Seidlitz et al., 2018; Ella & Keller, 2015; Nitzsche et al., 2018; McLaren et al., 2009; Quallo et al., 2010). Processing was performed with shell scripts using GNU parallel (Tange, 2011) for load balance. We used both affine transformations and non-linear warping (*3dQwarp*) (Cox & Glen, 2013) to generate four templates. Since the project was initiated before data collection was completed, 58 subjects from the first five cohorts were used to generate initial templates. The 12 subjects from the last cohort then were used as a validation set to test out-of-sample performance. After characterizing the 58-subject template we added these 12 to produce a full 70 subject template.

For simplicity of naming, we refer to the affine templates as ‘linear.’ Thus the four templates comprise 1) a 58 subject linear template (*T*_*L*58_), 2) a 58 subject non-linear template (*T*_*NL*58_), 3) a 70 subject linear template (*T*_*L*70_), and 4) a 70 subject non-linear template (*T*_*NL*70_).

To begin the procedure, each subject’s scan was AC-PC aligned in AFNI, which requires manual designation of the anterior commissure (AC), posterior commissure (PC) and the midsagittal plane. Image non-uniformity was corrected after manual skull-stripping. An iterative strategy was then used to produce successively refined templates by aligning each subject’s data to an existing template and then voxel-wise averaging all of the aligned data to form the next template. Transformation data sets for each subject were saved and later applied to selected landmarks for validation. To initiate the process, a single subject closest to the 58-subject median age and weight (5m, 17d and 24 kg) served as the initial template, *T* (0). Affine transformations aligned the remaining subjects to this one to produce *T* (1). After this, the linear and non-linear templates branched. Affine transformations produced *T* (2)_*L*_ and affine transformations followed by non-linear warping produced *T* (2)_*NL*_. Landmark validation errors served as the primary criteria for terminating the iteration. Based on this criteria, *T* (3)_*L*_ was not significantly better than *T* (2)_*L*_ and *T* (3)_*NL*_ was not significantly better than *T* (2)_*NL*_. Note that other quality assessments (tissue probability maps, spatial variance, and spatial signal-to-noise-ratio) had converged by this iteration as well. Thus our final *T*_*L*58_ and *T*_*NL*58_ corresponded to *T* (3)_*L*_ and *T* (3)_*NL*_, respectively. Finally to incorporate the remaining 12 subjects, we used the *T* (3)_*L*_ and *T* (3)_*NL*_ to align all 70 subjects. The voxel-wise averaging across all subjects then produced linear and non-linear templates that we refer to as *T*_*L*70_ and *T*_*NL*70_, respectively.

### 2.3. Characterization of the T_*L*58_ and T_*N*L58_ templates

#### Tissue Probability Maps

FSL (Smith et al., 2004) was used to generate tissue probability maps for each subject’s AC-PC aligned images. The *FSL-FAST* tool (Zhang et al., 2001) generated gray matter (GM), white matter (WM), and cerebrospinal fluid (CSF) maps. Subsequently, each subject’s respective affine and non-linear transformations were applied to the tissue maps. The voxel intensities were normalized from 0 to 1 within each tissue type and then averaged across all 58 subjects to create group CSF, GM, and WM tissue probability maps.

#### Contrast-to-Noise

We also used the GM and WM maps to calculate the contrast-to-noise ratio (CNR) between gray matter and white matter using the formulation in Nitzsche et al.(2015) (Nitzsche et al., 2015). Specifically, we calculated 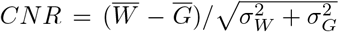 where 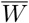, 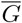, 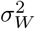, and 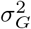 are the mean white matter intensities, mean gray matter intensities, the variance of the white matter intensities, and the variance of the gray matter intensities, respectively.

#### Spatial Characteristics

The spatial quality of the templates was assessed using measures of voxel-wise variance and signal-to-noise ratio (SNR), where SNR was computed as the mean voxel intensity divided by the voxel’s standard deviation across the 58 subjects. To quantify effective resolution of the templates we used the spatial Fourier transform to examine spectral power as a function of spatial frequency for both *T*_*L*58_ and *T*_*NL*58_.

#### Landmark Validation

Landmark validation used the AC, PC, and habenular nuclei (HB). The centroids of these locations were manually selected in the AC-PC aligned volumes for every subject as well as in the templates being evaluated. Most studies rely on the validity of a template by calculating RMS errors between fiducial landmarks within the same group of subjects that were used to create the template (Ella & Keller, 2015; McLaren et al., 2009; Conrad et al., 2014). We examined the internal error for *T*_*L*58_ and *T*_*NL*58_ by transforming each of the 58 subjects’ landmark coordinates to *T* (2)_*L*_ and *T* (2)_*NL*_ using their subject-specific transformations obtained from iteration 2 (recall that data transformed to *T* (2)_*L*_ and *T* (2)_*NL*_ were averaged to generate *T*_*L*58_ and *T*_*NL*58_). Template registration accuracy was then determined using the 12 out-of-template subjects from our final cohort as an unbiased measure. Affine registration accuracy was measured from the transformed subject’s landmark to the landmark in *T*_*L*58_ and *T*_*NL*58_ space as well as to the Göttingen minipig, *T*_*G*”_Γ*ottingen* (Watanabe et al., 2001)). To provide a fair comparison across the two Yucatan and one Göttingen templates, nonlinear registration would have confounded the warping algorithm’s ability to compensate for inter-breed distortions.

## 3. Results

Figure 1 shows the final (*T*_*L*58_) and (*T*_*NL*58_) templates. For a visual comparison, the top row of Fig. 1 shows the median subject that served as the initial registration target, *T* (0), and is thus also representative of the data collected for the cohort. The axial, sagittal, and coronal views were defined by the templates’ origin at the anterior commisure. A visual comparison demonstrates that the non-linear template has enhanced outer edge boundaries and greater anatomical detail surrounding WM, GM, and ventricles compared to the linear template. These differences are perhaps best highlighted by comparing the coronal slices. In addition to visual inspection, we used several measures to characterize the quality of these templates.

**Figure 1:**
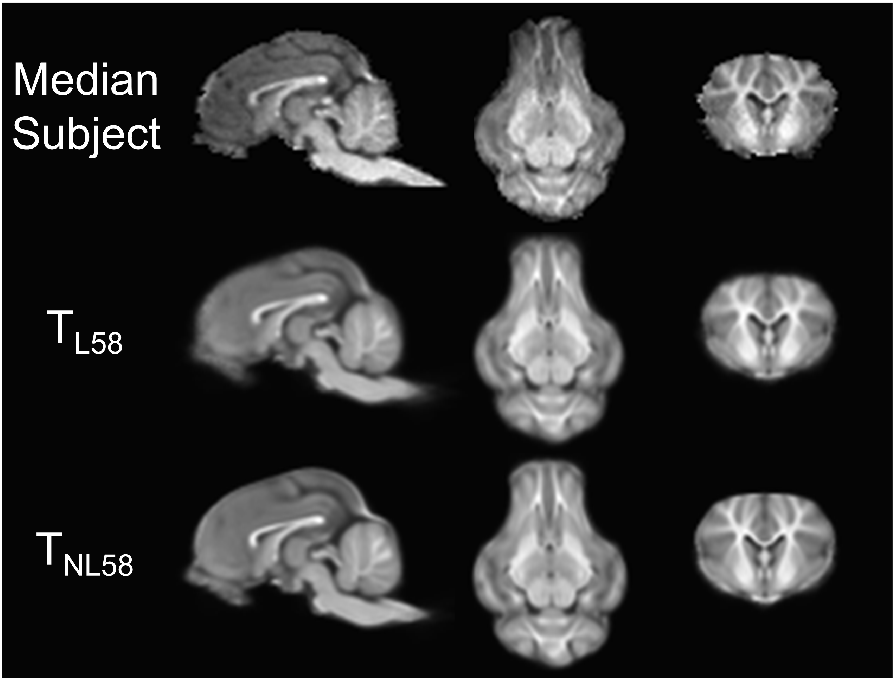
The final (*T*_*L*58_) and (*T*_*NL*58_) templates. Shown are the sagittal, axial, and coronal views of the median subject (5m, 17d and 24 kg) compared to the final 58 subject linear and non-linear templates. In the top row, the median subject is skull-stripped, intensity-corrected, and AC-PC aligned. In all three volumes, the horizontal axis is set to the AC-PC line and the origin is set at the anterior commissure (AC).

### 3.1. Tissue probability maps

Figure 2 shows the tissue probability mapping results. Figure 2A demonstrates that both *T*_*L*58_ and *T*_*NL*58_ have sufficient contrast to create high probability masks of CSF, GM, and WM. By varying the probability threshold equally across these maps, we generated a plot of CNR between gray matter and white matter vs. tissue probability threshold (Fig. 2B). Based on visual comparison of Fig. 2A and the CNR characteristics of Fig. 2B, both templates are highly similar. However, note that the CNR for the *T*_*NL*58_ is greater than in the *T*_*L*58_ for thresholds below 0.5 and then flips so that *T*_*L*58_ is significantly greater at the 0.9 threshold (*p* < 0.05).

**Figure 2:**
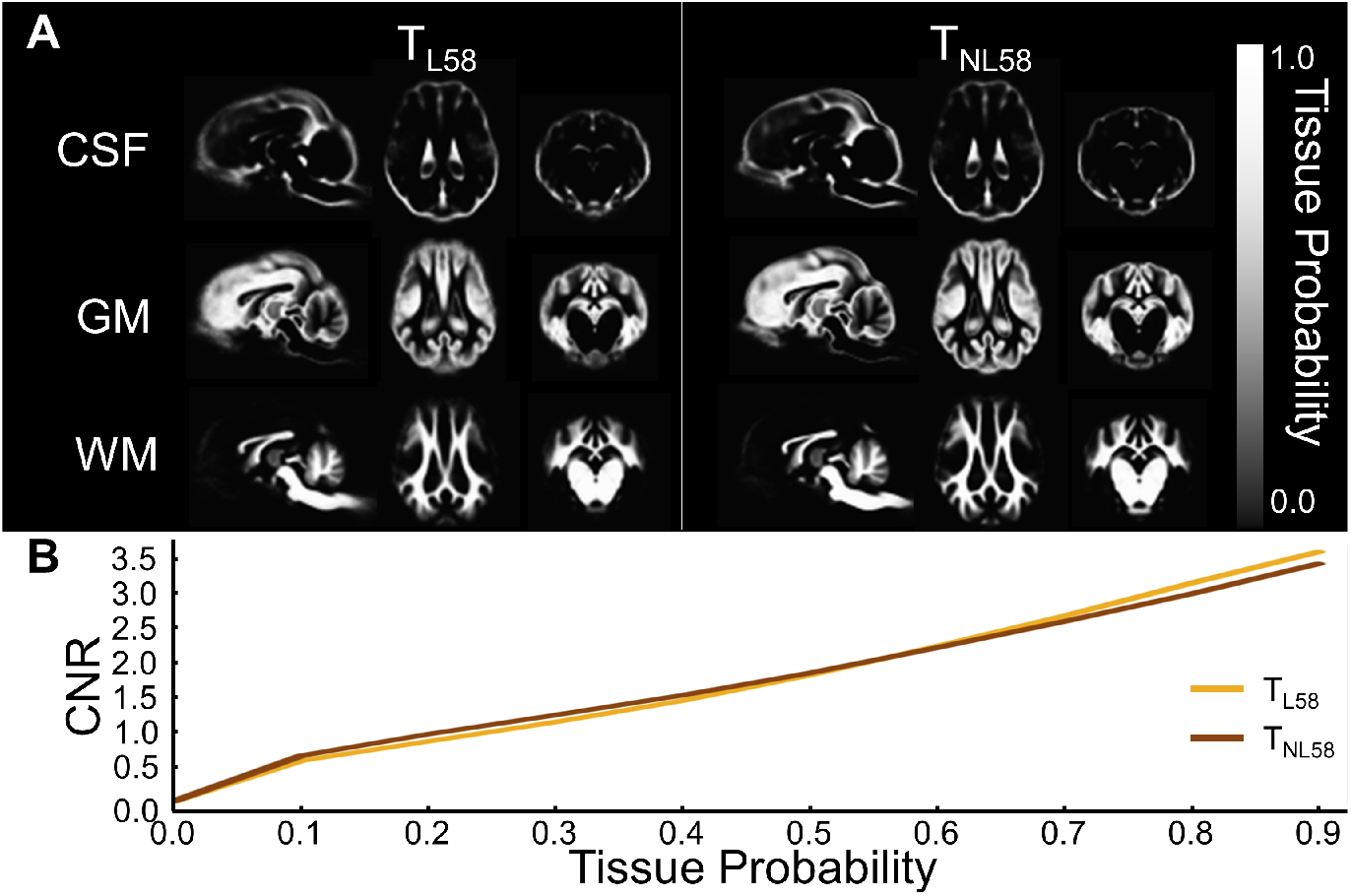
Tissue Probability Maps. (A) CSF, GM, and WM probability maps for *T*_*L*58_ and *T*_*NL*58_. (B) Whole-brain gray matter and white matter CNR as a function of tissue probability threshold.

### 3.2. Spatial characteristics

Figure 3 maps the voxel-by-voxel variance and SNR for the templates. In both cases, these metrics were calculated across all subjects after alignment to *T* (2)_*L*_ and *T* (2)_*NL*_. Recall that averaging these volumes generated *T*_*L*58_ and *T*_*NL*58_. Figure 3A shows the spatial variance at each voxel. For display purposes, the variance was normalized by the maximum observed between the two maps. As shown, the regions with high variance include the edges of the brain, the olfactory bulb, and brain stem. Visual inspection demonstrates that the general pattern of variance is similar but reduced for *T*_*NL*58_. Figure 3B complements Figure 3A, but the two metrics are not redundant. Unlike the variance maps, the SNR is lowest at the edges of the brain and higher for internal structures. The highest SNR is found in the white matter for both templates. Comparing across linear and non-linear templates, the non-linear again shows improvement. In this case, *T*_*NL*58_ has an increased distribution of high SNR.

**Figure 3:**
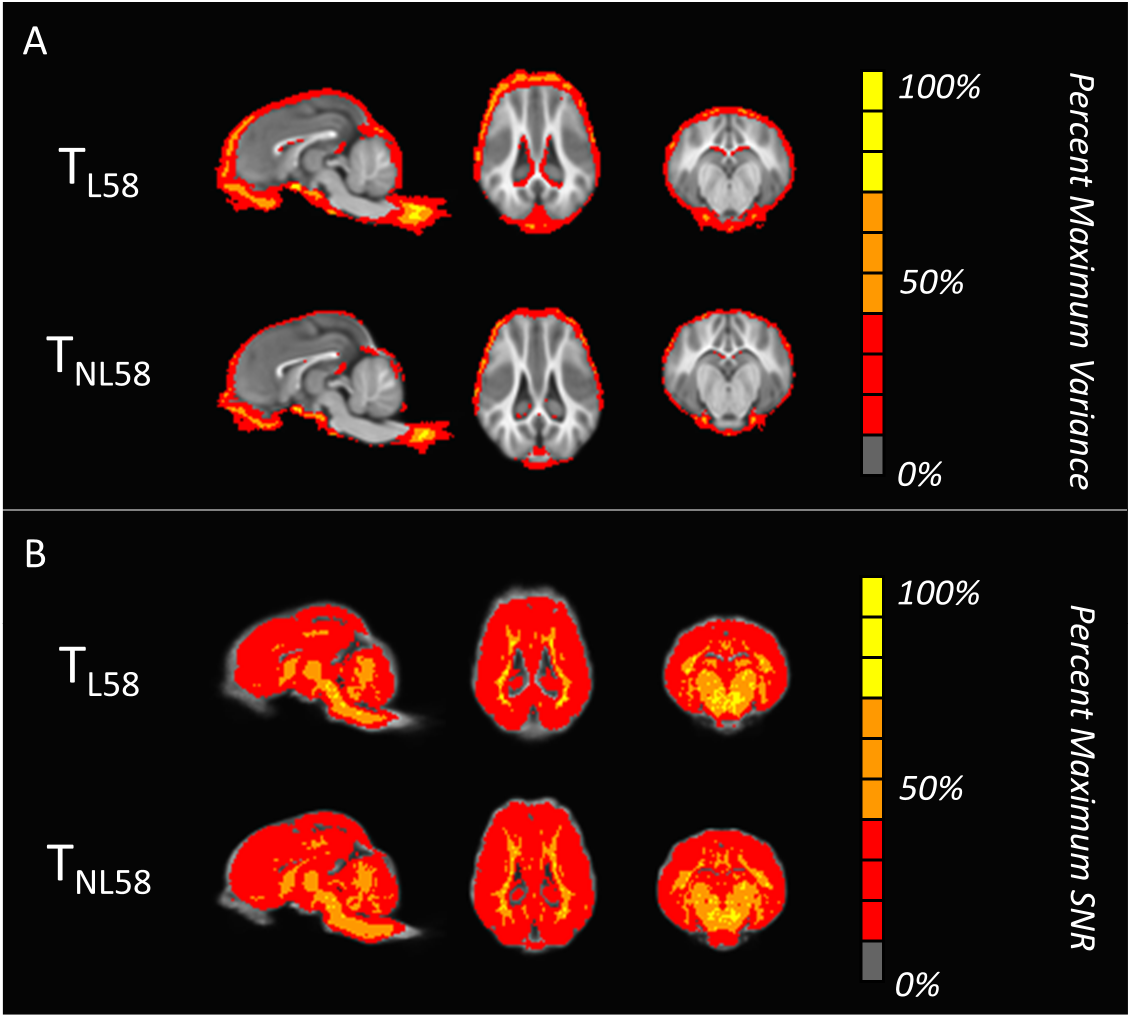
Quality Inspection. (A) Undesirable spatial variance is concentrated along the edges of the template at a greater magnitude and extent in *T*_*L*58_. (B) Desirable SNR is greater within the internal structures and is enhanced in *T*_*NL*58_.

Figure 1 qualitatively suggests that the nonlinear template has a higher resolution than the linear one. This is characterized further in Fig. 4, which evaluates the effective spatial resolution for the median subject and compares this to *T*_*L*58_ and *T*_*NL*58_ using the spatial Fourier transform. Fig. 4A shows the axial slices that were used for this analysis. The magnitude Fourier transforms of the slices are shown in Fig 4B. Even though both templates as well as each subjects’ data have a nominal 1 *mm*^3^ isotropic resolution, the spatial Fourier transform provides a quantitative comparison of the effective spatial resolution of these three images. The center of the spectra represent the mean of each image. Moving outward from the center represents the proportion of image power contained at higher spatial frequencies. Thus higher magnitudes at higher spatial frequencies generally indicates better effective resolution. The major caveat to this is that noise tends to augment the magnitude at all frequencies. For example, Gaussian white noise distributes uniformly across all frequencies and thus creates a flat noise floor. For these images, the highest spatial frequency is Nyquist-limited to 1/2*mm*^−1^ in each dimension. To simplify comparisons, Fig. 4C plots the profiles for the spectra along the colored lines indicated in Fig. 4B. Figure 4C displays an increase in quality in *T*_*NL*58_ compared to *T*_*L*58_. Not surprisingly, however, effective resolution is reduced in both templates compared to the median subject, which demonstrated the largest magnitudes at higher spatial frequencies. The templates’ reduced effective resolution is due to heterogeneity across subjects as well as registration and interpolation errors. Note, however, that both templates preserve a substantial amount of effective resolution while averaging over a large number of subjects. Moreover, it is important to note that the median subject (and all other individual subjects not shown) also has a grainy appearance and certainly consists of some level of noise leading to high spatial variation even across uniform tissue types. Thus the decreasing magnitude in the templates at higher frequencies also demonstrates that the averaging across volumes to generate the templates acts as a low pass filter. Arguably, the non-linear alignment provides a more specialized filter that preserves power at higher spatial frequencies and therefore preserves edges and boundaries, while reducing high spatial variation that leads to image graininess.

**Figure 4:**
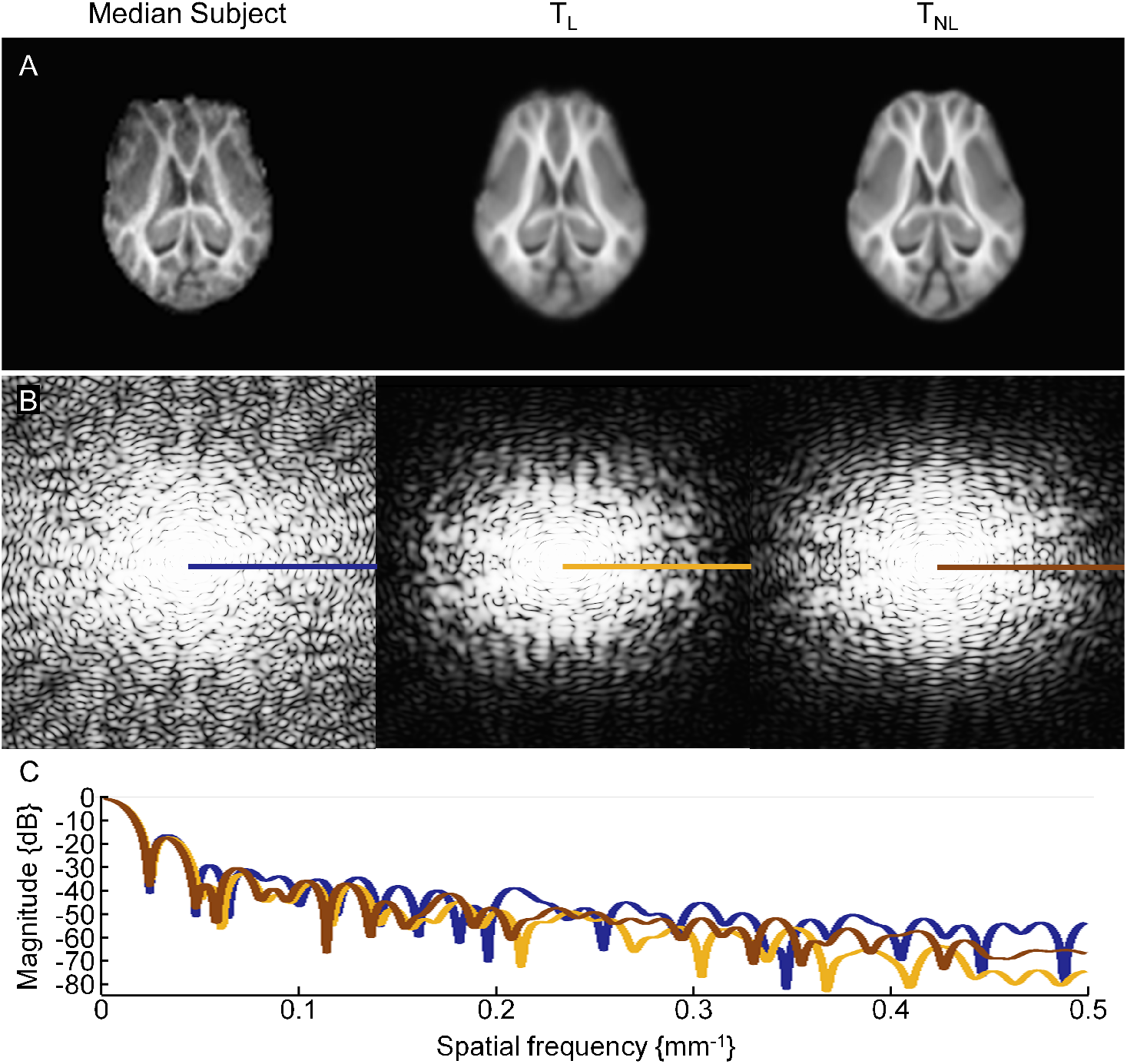
Effective spatial resolution. The 2D spatial Fourier transform was applied to the data shown in (A) to produce the magnitude spectra displayed in (B). The plots in (C) are representative spectra taken from the colored profiles indicated in (B). For visual comparison, each of these spectra were normalized by their respective maximum magnitudes.

### 3.3. Landmark Errors and Alternative Template

The internal validity of the *T*_*L*58_ and *T*_*NL*58_ templates is quantified in Table 1, which lists the average and maximum distances for the AC, PC, and HB across the 58 subjects. On average, the landmark errors were approximately one voxel in linear dimension (1 mm), with the greatest errors found in the AC. In addition, the final 12 subject cohort was used as an independent (out-of-sample) validation set to compare registration accuracy to *T*_*L*58_ and *T*_*NL*58_ and the only alternative Göttingen minipig template (Watana_b_e et al., 2001). See Table 2. The out-of-subject performance between the *T*_*L*58_ and *T*_*NL*58_ showed no significant difference, indicating that the warping transformations did not affect the overall registration accuracy. However, compared to the Göttingen template, both templates had significantly improved registration accuracy in the PC (*F* = 18.27, *p* < 0.0001) and the HB (*F* = 10.77, *p* = 0.0003).

**Table 1:** Average and maximum distances from the template landmarks (in mm) between the 58 subjects and the *T*_*L*58_ and *T*_*NL*58_ templates.

|  | Anterior<br>Commissure |  | Posterior<br>Commissure |  | Habenuar<br>Nuclei |  |
| --- | --- | --- | --- | --- | --- | --- |
| Template | Mean | Max | Mean | Max | Mean | Max |
| $T_{L58}$ | 1.00 | 2.11 | 0.82 | 1.54 | 0.79 | 1.51 |
| $T_{NL58}$ | 1.03 | 2.07 | 0.79 | 1.44 | 0.79 | 1.48 |

**Table 2:** Average and maximum distances from the centroid of the template landmarks (in mm) following registration of the 12 validation subjects to the *T*_*L*58_, *T*_*NL*58_, and Göttingen templates.

|  | Anterior<br>Commissure |  | Posterior<br>Commissure |  | Habenuar<br>Nuclei |  |
| --- | --- | --- | --- | --- | --- | --- |
| Template | Mean | Max | Mean | Max | Mean | Max |
| $T_{L58}$ | 0.82 | 1.34 | 0.84 | 1.56 | 0.81 | 1.05 |
| $T_{NL58}$ | 0.99 | 1.75 | 0.82 | 1.69 | 0.89 | 1.19 |
| $T_{G\ddot{ot}tingen}$ | 1.08 | 2.06 | 1.52 | 2.29 | 1.23 | 1.75 |

## 4. Discussion

A multi-modal TBI study created an opportunity to produce a highly specialized template of the five- to seven-month-old male Yucatan minipig based on 70 subjects. Our initial objective was to apply non-linear warping to develop a T1-weighted template of the male Yucatan minipig using 58 subjects. Before finalizing this template, however, we had collected an additional 12-subject cohort. We also decided early in the process that producing a linear (affine) template required only minimal additional effort while providing an opportunity to make comparisons with the non-linear one. Thus, we have produced and archived four templates that are suitable for use in neuroimaging analysis pipelines. We are not aware of previous template characterizations that have an additional “left-out” validation set, and our approach raises at least two issues. The first is that this suggests the possibility of doing a fully cross-validated study. We decided not to pursue this because of the high computational costs, and the additional burden of interpretation. More importantly, though, because this data set is so homogeneous across a number of dimensions (same scanner, all males, narrow age range, etc.) and based on the uniformity/consistency observed between the first 58 subjects and the final 12 subjects in Tables 1 and 2, the likely utility of a fully cross-validated study is low in this case. We note, though, that a fully cross-validated approach maybe worth further investigation and would likely complement consensus-based template creation processes such as the one proposed by Avants et al. (2010), which aim to produce templates that are not biased by any one of the template subjects. The second issue is that the use of the added validation set implies that the 58-subject templates are more fully characterized than the *T*_*L*70_ and *T*_*NL*70_. But based on the minor differences that we have observed, we expect that the quality of the 70 subject templates are equivalent or better than *T*_*L*58_ and *T*_*NL*58_. In addition, we anticipate marginally enhanced performance from the non-linear templates. While relatively comparable, the non-linear template was equal to or slightly better than the linear template in all qualitative and quantitative assessments, with the exception of the small but statistically significantly greater gray-white matter CNR in *T*_*L*58_ occurring at very high tissue probability levels. Thus, our recommendation is to use the 70 subject non-linear template (*T*_*NL*70_.). Practically, however, we believe that all four templates will achieve similarly acceptable results.

Wilke et al. (2017) has thoughtfully summarized many of the issues and trade-offs that occur along the continuum of registrations that span low dimensional affine transformations to high dimensional warping. While we used rather conventional techniques, the simultaneous generation of both affine and non-linear templates is unusual. At a basic level, this allowed us, to evaluate the quality and relative merits of both approaches during the actual process of template generation. Visually, it is not surprising that the non-linear looked better. In fact, this will always be the case since the aim of warping is to decrease residual group variance (Wilke et al., 2017). By then looking at spatially localized measures of variance and SNR across subjects as in Fig. 3 and by also examining globally diagnostic measurements using the Fourier transform to examine effective resolution in Fig. 4, we see modest advantages to the non-linear warping approach. Indeed, our homogeneous cohort probably presents a special case. The similarity between *T*_*L*58_) and *T*_*NL*58_ is likely strong confirmation that this is a highly uniform sample of subjects.

Beyond qualitative visual assessments of Fig. 1, the quality of the templates was quantified by tissue segmentation (Fig. 2), voxel-wise variance and SNR (Fig. 3), effective resolution (Fig. 4), and landmark errors (Tables 1 and 2). Tissue segmentation worked well in both the linear and non-linear Yucatan templates. One issue that has been historically troublesome in non-human templates has been the lack of contrast (Seidlitz et al., 2018). Visually, Fig. 2A demonstrates that that the templates have sufficient contrast to enable gray, white, and CSF mapping. As noted, the tissue probability maps are sharper for the *T*_*NL*58_. Although both templates were similar across all tissue probability levels, CNR was higher across subjects for *T*_*L*58_ at the 90% threshold. The CNR for both *T*_*L*58_) and *T*_*NL*58_ is high relative to other estimates reported in the literature. Specifically, they are in the same range as (Nitzsche et al., 2015) who reported a CNR of approximately 1.85 for a non-linear sheep atlas. Both *T*_*L*58_ and *T*_*NL*58_ exceed this level beyond the 70% probability threshold. We note also that the objective here was to produce templates for neuroimaging processing pipelines for functional and structural analyses, and we have not considered their utility for surgical planning, although subcortical contrast is also a major factor for surgical planning in experimental procedures (Rosendal et al., 2010).

The maps of voxel-wise variance and SNR measures highlight the individual brain structure variability and should also be diagnostic of alignment accuracy. The most noticeable features of the variance maps in Fig. 3A show that the outer surface of the brain, the edges of the ventricles, the olfactory bulb, the brainstem, and the transverse fissure separating the cerebellum from the cerebrum varied most prominently across subjects. Notably, the variance decreased for the *T*_*NL*58_ compared to the *T*_*L*58_. Still, voxel-wise variance in the non-linear template remains prominent in the brainstem and olfactory bulb and residually present in the ventricles, suggesting that these are regions of relatively higher inter-subject variability. The variance at the outer edges also reflects variability in brain volume across each subject. However, this surface variance also likely reflects imperfect skull-stripping. Nonetheless, there are no obvious systematic errors in the skull stripping and the large number of subjects resulted in templates with smooth, crisp edges that plausibly represents the subject population. The brainstem includes a possible third source of variability. In addition to anatomical variability and manual skull stripping, the high variance also likely arises from variation in neck angle during scanning. The voxel-wise SNR estimates of Fig. 3B complement the variance maps. The most striking shared properties across the two metrics are the relatively low SNR (high variance) at the edges of the brain and the higher SNR (lower variance) for internal structures. Like the variance maps, SNR improves in the *T*_*NL*58_. While variance highlighted regional inter-subject variability, the SNR maps highlight similarities. For example, the white matter is most prominent in the *T*_*NL*58_ SNR maps, indicating structures that are highly conserved across the study subjects.

The Fourier analysis in Fig. 4 suggests that the templates preserve spatial resolution while also filtering some noise from the original data. In general contexts, signal has finite support while noise spans the entire Fourier space. Our analyses suggest that the attenuation in higher spatial frequencies of the templates compared to the median subject appear to be a reasonable balance between filtering noise and preserving spatial signal. Based on the axial slices shown in Fig. 4, we would expect most of the power to be concentrated in an oval - since the image data are longer from anterior to posterior, this direction has a longer period (lower spatial frequency) compared to the left to right direction. This oval relationship is less pronounced in the median subject’s power spectrum compared to the two templates. In addition, it is desirable to preserve as much high frequency signal as possible, since this reflects the templates’ effective spatial resolution. Inspection of the tails in Fig. 4C shows that the non-linear filter has preserved power at a level that is approximately the mid-level between the original data and the linear template.

Distance variations for the AC, PC and HB landmarks were calculated within the 58 subject templates and are reported in Table 1. The AC and PC have been used almost ubiquitously in past studies (Conrad et al., 2014; Ella & Keller, 2015; McLaren et al., 2009; Black et al., 1997, 2001; Love et al., 2016). The HB was selected here as a third measure that was independent of the AC-PC alignment and could serve as a distinct location that could be consistently manually labeled. The HB followed similar trends to the AC and PC, however, it is expected that landmarks selected along the outer regions of the brain would have diminished registration accuracy due to the increased template variance along the edges. Indeed, as pointed out by (Rohlfing, 2012) for non-linear methods, only a very dense set of landmarks can fully characterize registration accuracy. In previous reports, the neonatal piglet template (n=15) showed a mean variation of 0.41 and 0.65 mm (maximum 0.72 and 1.07 mm) for the AC and PC landmarks, respectively (Conrad et al., 2014). In a sheep brain template (n=18), the average distance from AC and PC points was about 0.44 and 0.56 mm, respectively, with maximum distances of 1.0 and 1.2 mm (Ella & Keller, 2015). Similarly, the rhesus macaque template (n=82) had an average variation of 0.8 and 0.8 mm with maximum distances of 1.87 and 2.24 mm (McLaren et al., 2009). Our numbers in Table 1 are thus relatively high. Interestingly, though, this effect seems to be closely correlated with the large voxels used here. We acquired at 1 × 1 × 1 *mm*^3^, which corresponds to a cubic voxel dimension of 1 mm. Conrad et al. (Conrad et al., 2014) had 0.35 × 0.35 × 1 *mm*^3^ (corresponding to an effective voxel length of 0.64 mm). Ella et al. (Ella & Keller, 2015) used 0.5 × 0.5 × 0.5 *mm*^3^ (0.5 mm voxel length) and McLaren et al. (McLaren et al., 2009) used data from several sites, but had an effective linear resolution of approximately 0.57 mm. When normalizing fiducial distance errors by these effective cubic voxel sizes, our Yucatan templates as well as those of Conrad et al. (Conrad et al., 2014) and Ella et al. (Ella & Keller, 2015) have an approximately 1:1 ratio, while the McLaren et al. (McLaren et al., 2009) results produce a factor of approximately 1.4. Thus it appears that most non-human templates have errors that compare closely with their effective cubic voxel size, which is reasonable since this is the major limiting factor for defining the centroids of fiducial markers. Based on this, the relatively low resolution of our T1-weighted data is its major limitation and should be the primary factor to target for improving the quality of future templates.

The 12-subject validation set provided an independent measure of landmark variations to assess internal bias that could arise from the standard practice of using the same subjects to both create and test a template. As mentioned, when comparing results between *T*_*L*58_ and *T*_*NL*58_, these twelve had less variation in terms of mean and max than the internal 58-subject results. This is the opposite of what would be expected if within-sample bias were present. Since any internal bias from creating the template and then testing with those same subjects should have made Table 1’s results appear better, the out-of-sample validation provides strong evidence that such bias is not a factor in the 58 subject templates. We also used these 12 subjects to assess the utility of the 0.473 × 0.473 × 1.125 *mm*^3^ Göttingen template (Watanabe et al., 2001)). Even though, the nominal voxel size was smaller for the Göttingen template, the registration errors were higher for the 12-subject validation set compared to both *T*_*L*58_ and *T*_*NL*58_. This confirms the utility of specialized templates.

## 5. Conclusion

While the minipig is growing in experimental popularity, there is currently a lack of appropriate brain templates for neuroimaging pipelines to support structural and functional studies. We have generated and compared linear (affine) and non-linear templates in a large, homogeneous population of Yucatan minipigs. We have also validated templates from 58 subjects using an additional 12 subject validation set. Our characterization of these templates across visual appearance, spatial SNR, gray-white matter CNR, effective resolution, and landmark coordinate variation generally found that the non-linear approach was slightly better than the linear one, and that there was no strong evidence for internal bias of the 58 subject templates. Both of these factors are most likely due to the uniformity of the subject population. All original and AC-PC aligned skull-stripped data, processing scripts, and four resulting templates, *T*_*L*58_, *T*_*NL*58_, *T*_*L*70_, *T*_*NL*70_, are archived at [https://lacontelab.github.io/VT-Yucatan-MRI-Template/]. While only minor differences have been noted, we recommend *T*_*NL*70_ for future use.

## 6.Acknowledgements

The authors would like to thank Christopher Anzalone, Douglas Chan, Shaylen Greenberg, and Allison Guettler for their assistance with image acquisition, Alana Hull for comments, Daniel Glen and Robert Cox for comments and support. MRI scans provided by Office of Naval Research award N0001414-C-0254.

## Notes

https://lacontelab.github.io/VT-Yucatan-MRI-Template/

### Competing Interest Statement

The authors have declared no competing interest.

